# Discovery Function Labels Based On Numerical Features Of MicroRNA Found In Plant

**DOI:** 10.1101/770875

**Authors:** Rongsheng Zhu, Jinglu Liu, Dawei Xin, Zhanguo Zhang, Zhenbang Hu, Yang Li, Qingshan Chen

## Abstract

MicroRNAs(miRNAs),are a class of small endogenous non-coding RNAs that play Important post-transcriptional regulation role by degrading targeted mRNAs or repressing mRNA translation. We screened 84 miRNAs belonging to 21 conserved family from 4014 miRNAs in miRBase database which distributed in 47 plant species. Of the predicted 274 target genes,42 GO terms were found in the Gene Ontology. With the 135 numerical features which extracted by perl program, the difference significantly result of ANOVA (P<0.001) and multiple comparison show that the function labels G…,GC content and Helix in biological process, cellular component and biological function, respectively. Our result have suggested a potential connection between numerical features of miRNAs and the function of target genes of miRNAs.

## 1. Introduction

MicroRNAs(miRNAs),are a class of small endogenous non-coding RNAs that play Important post-transcriptional regulation role and found in plant and animal. Plant miRNAs, with a mature length of approximately 22nt, recognized nearly perfect complementary binding to 3’-UTRs of target mRNA[1–3]. Recent studies have suggested that miRNA have essential function including developmental timing, cellular differentiation, proliferation, apoptosis, cell identity and fate and response to environmental stress[1, 4–8]. Since the first miRNA lin-7 and lin-4 was found in C. elegans have been more than two decade[9],the previous“Junk DNA”is considered to play an important role in the biological processes.

With the development of computational identification[10, 11], majority of the new function of predicted miRNAs and their numerical features have been found. For example, previous studies have shown that unlike transfer RNAs and ribosomal RNAs, the minimum free energy(MFE) of majority miRNAs are lower than shuffled sequences[12]. The adjusted minimum free energy (AMFE)[13], and the minimum free energy index(MFEI)[14] also indicate that the high tendency of stable secondary structure of miRNAs. Although the small mature miRNAs segment with high degree of similarity across species,however, it is difficult for experimental verified and high cost[15]. Computational analysis of numerical features of miRNAs is a new research method that may bring new discoveries for the miRNAs research with a new vision.

*A.thaliana*, a model crop, as a major material in biological research which has a small genome, low repetitive sequences and easily converted. Previous studies suggest that miR156 could inhibit transcripts of zygotic SPL as to prevent premature accumulation in embryonic maturation phase in *A.thaliana* [16]. The miR395 family which was predicted to code for ATP sulphurylases by target mRNAs[1, 17]. The miR162 and miR168 regulate genes required for miRNA biogenesis or activity [18, 19]. All of these revealed that functional diversification of conserved miRNA which also highlighted the importance of research in numerical features. Because the numerical features reflect the structure and function significantly[20]. Whether these miRNAs with different function related to the differences of numerical feature? In this paper we aim to try to find the numerical features to explain the diversity of functions.

In this study, we used ANOVA and Multiple comparison to evaluate significant differences between numerical features from 21 conserved families and GO terms of predicted targets. The conclusions as follow:first, We got 274 predicted target genes and found that they enriched in 42 GO terms which 23 in biological process, 11 in cellular component and 8 in molecular function. Second, as the result of ANOVA, there are significant difference between numerical features and the GO terms, respectively. Third, we got function labels based on numerical features of miRNAs in plant. The secondary structure feature G…, GC content and Helix show significant difference in biological process, cellular component and biological function, respectively. Our conclusions reveal the relationship between numerical features of miRNAs and its functions.

## 2. Materials and methods

### 2.1 Data prepare

#### 2.1.1 MiRNA genes and predicted target

We selected 4014 miRNA genes from miRBase Sequence Database, release 18 (http://www.mirbase.org/)[21]which covered 47 species in viridiplantae. We screened 84 miRNA genes belong to 21 conserved families in plant. All 274 predicted targets genes in *A. thaliana*, obtained from PMRD (http://bioinformatics.cau.edu.cn/PMRD/)[22]. Detailed information is listed in supplemental Table 1.

### 2.2 Data analysis

#### 2.2.1 Numerical features of miRNA

We designed a perl program for assessing 135 numerical features from the selected miRNA genes. These features were roughly divided into these classes which contain frequency characteristics of nucleotides, frequency characteristics of the secondary structure matching state, energy features and entropy features. details see our previous study by Zhu et al [23]. The novel numerical features of external loop, hairpin loop and multi-loop are added in this study. (see Table 2)

#### 2.2.2 Go annotation

In order to detect functional categories enriched in genes targeted by selected conserved miRNA family, we analyzed the annotation of transcripts with putative target sites in The Gene Ontology(http://www.geneontology.org/, AmiGO version 1.8)[24]. Matlab was used to visualize the enriched the GO terms and miRNAs.(see Fig. 1)

**Fig. 1.**
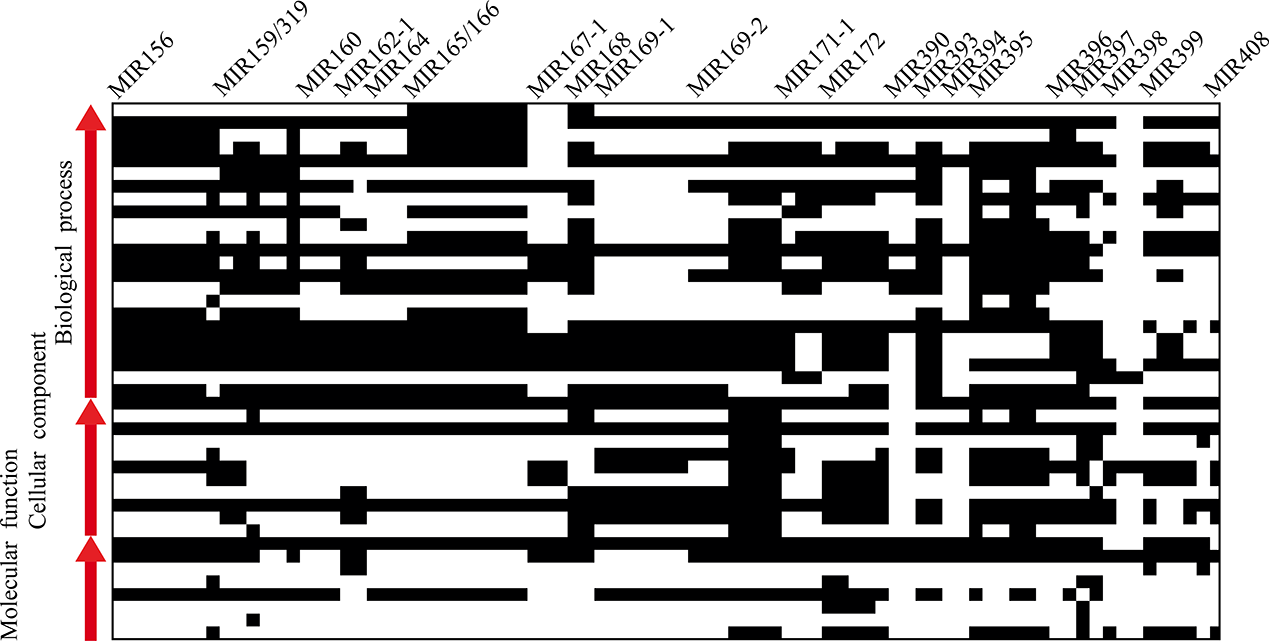
Cluster GO terms and conserved miRNAs. The x-axis represents 84 conserved miRNAs, the y-axis represents the 42 GO terms in which the three GO classification. The number of biological processes, cellular component and molecular function is 23,11 and 8, respectively. The white box represents present and black box represents absence. The analysis results are summarized in table

#### 2.2.3 Basic statistical methods

We used ANOVA (P<0.001)to evaluate significant differences in biological process, molecular function and cellular component, respectively. Multiple comparison was used to further analysis the significant result of ANOVA.

## 3. Result

### 3.1 Screening conserved miRNA families

We got 21 conserved miRNA families from 4014 plant miRNAs in miRBase database which coverd 47 plant species. The miR156 is the most conserved miRNA families which coverd 34 plant species, and the relatively conserved miRNA families miR162-1/394 coverd 14 plant species. Most of the conserved families are multigene families in which each miRNA product highly similar mature miRNAs. In general, more conserved miRNAs have a tendency to express at a higher levels than do relatively conserved miRNAs[25]. In the case of miR166 binding the mRNAs which encoded class III HD-ZIP transcription factors, the complementary sites are almost identical in seed plants, lycopods, and mosses, hornworts and liverworts[26, 27]. It suggested that the conserved miRNA families tend to regulate the same function in different species. (seetable.1)

### 3.2 Functional diversification of predicted target genes of miRNAs

In this study,we focused on the relationship between GO terms and conserved miRNAs. We mapped the predicted targets genes of conserved miRNAs to the Gene Ontrology (Fig.1).we find that the predicted miRNA target genes from conserved families have a significant enrichment. The result showed that there were more than 75 miRNAs enriched in 9 GO terms for each. The GO term of ‘Metabolic process’ which enriched the most miRNAs. On the contrary, there are only 4 miRNAs enrich in ‘Pigmentation’, 5 in ‘Electron carrier activity’, ‘Enzyme regulator activity’ and ‘Receptor activity’. Only 2 miRNAs enriched in ‘Structural molecule’. These results suggested that conserved miRNA exercise with a variety of regulatory functions in the process of life activities. As shown in Fig.1. We also detected that the members of conserved miRNA families of miR160, miR164, miR165, miR167-1, miR169-1 which have the same enrichment, respectively. There are only 9 and 8 GO trems in miR390 and miR394. This result indicated that conserved mature products of miRNAs of plant have a tendency to function consistency. Interestingly, we find that different member of conserved miRNAs familes have different function category, such as miR-159(319)、miR-171-1、miR-395 and miR-397.

### 3.3 Significant difference among GO terms based on numerical features of miRNAs

In order to investigate whether there is a significant diference between GO terms and 135 numerical features, we used ANOVA to analyze the differences between them (Fig.2). The ANOVA (P<0.001) results showed significant differences in these three GO categories. As shown in Fig. 2A, the numerical features A%, G…,stack,(G+C)%,TE (information entropy of triple nucleotides) in biological process showed significant difference with their sequence number of 1, 93, 123, 128, 134. It is noteworthy that the five of numerical features in cellular component GC content, GCG, UGG, G‥+, (G+C)/(A+U) belong to the sequence number 14, 42, 78, 102, 129 show in Fig.2B, which are all closely related to nucleotide frequent, secondary structure and information entropy feature. We also found some strong difference in Fig.2C, the numerical feature of Bulge loop, Helix, GA, AGC and SSE (Secondary structure paired state information entropy) may be important in the molecular function.

**Fig. 2.**
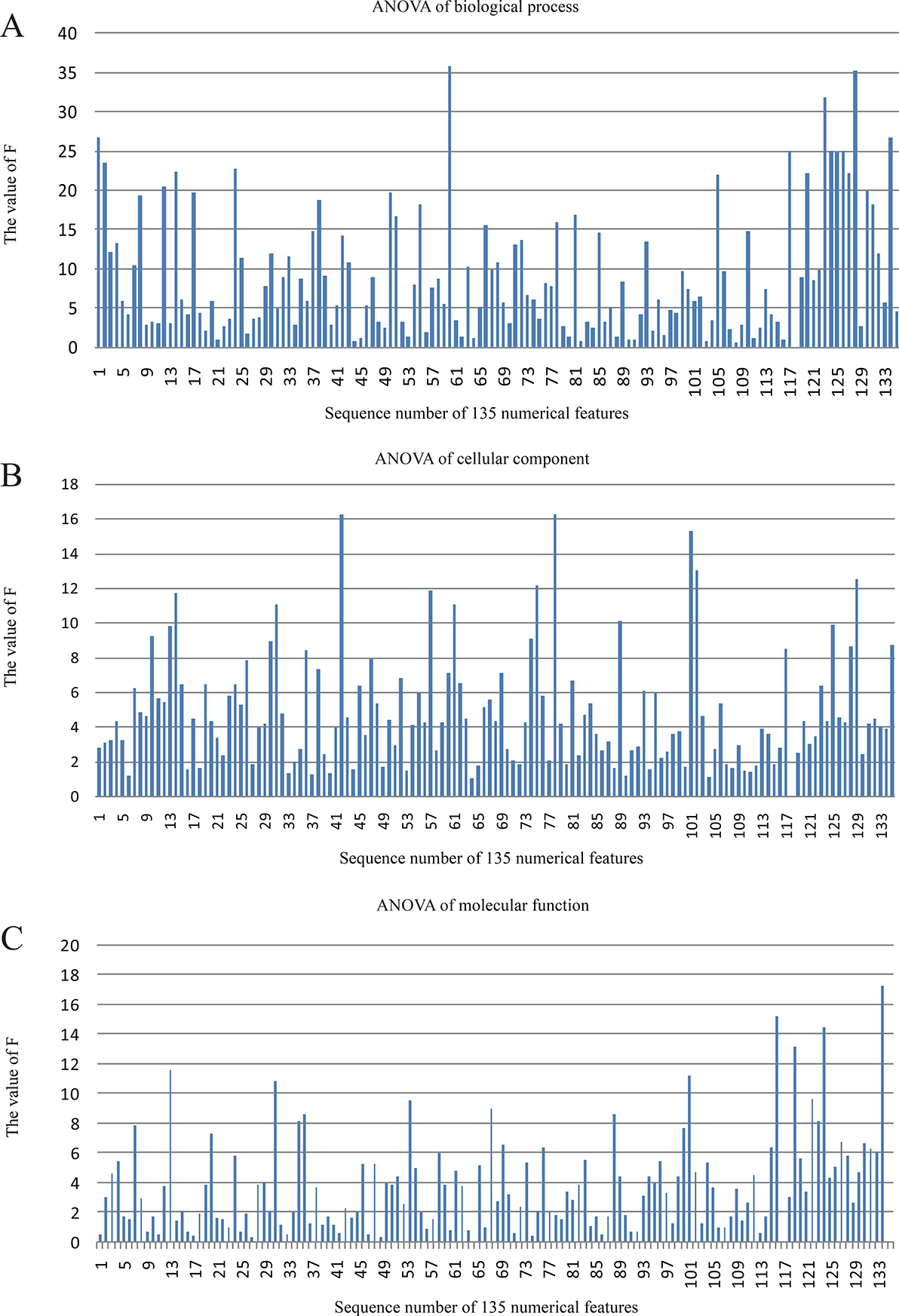
The analysis indicated the relationship between numerical features of miRNAs and GO terms at GO categories level. A, B and C show the result of ANOVA based on three GO classification and 135 miRNA numerical features. We used the F value to evaluate the difference. Original data have been standardized in table2

### 3.4 Function labels based on numerical features of miRNA

Whether there is a class of numerical features which could distinguish between different biological function of the target gene in these significant difference numerical features? With this question, we extended a further analysis, based on the result of ANOVA, Multiple Comparisons used to the results of significant difference selected from ANOVA(P<0.001). We find some function labels which could distinguish the function at numerical features level. The result indicated that the result of Multiple Comparisons in three GO categories all have a significant different numerical feature.

The probability density curve and the case of structure of miRNAs (Fig.4 A, D) show that miR398a/172e could distinguish the GO term ‘establishment of localization’ and ‘metabolic process’ at the numerical feature G… in biological process. Fig.4 B, E show that miR398b/160a could distinguish the GO term ‘cell’ and ‘membrane’ at the the numerical feature of GC content. Fig.4C.F show that miR162a/164a could distinguish the GO term ‘catalyic activity’ and ‘nucleic acid binding transcription factor activity’. The secondary structure feature G…, Helix and GC content with significant result indicated that these three features can be used as the specific label on the miRNA level to distinguish the biological functions.

## 4. Discussion

Our analysis showed that the highly conserved miRNAs families have a tendency to the regulation of target genes consistently (Fig.1) The different members of conserved miRNAs families tend to enrich in the same terms. Owing to the highly conserved mature of miRNAs and the extensive complementarity with target genes, indicating to regulate the same target genes and to exercise the same function. The result also show that the same miRNA enrichment in multiple functional classification, suggestting the functional diversity of miRNA regulation. Previous studies have shown that microRNA families were conserved between plant species and target sites were also conserved in target mRNAs[25, 29–31]. We used the measure of numerical features to evaluate conserved miRNA sequences. Because of the numerical features of miRNA reflect the structure and function.

In order to evaluate the numerical features of conserved miRNAs whether there were a Intrinsically linked to the biological activities. Analyzes using ANOVA (P<0.001) at GO terms level(Fig.2) showed that there are some significant difference numerical features. The result indicated that these shared significant difference numerical features is likely related to the biological activities of miRNA target genes(Fig.3). However, those differences were not significant one is likely to be a stable part of supporting biological activities. The multiple comparisons result suggested that the numerical feature of miRNA could distinguish the GO terms. The numerical feature G… wsa the function lable of ‘establishment of localization’ and ‘metabolic process’ in biological process. Previous studies have shown that miRNA398 likely regulates plant responses and the plant stress regulatory network, like oxidative stress, water deficit, salt stress, abscisic acid stress and so on [32]. This feature likely to be tend to affect the secondary structure of the pre-miRNA, often associated with an internal loop structure.

**Fig. 3.**
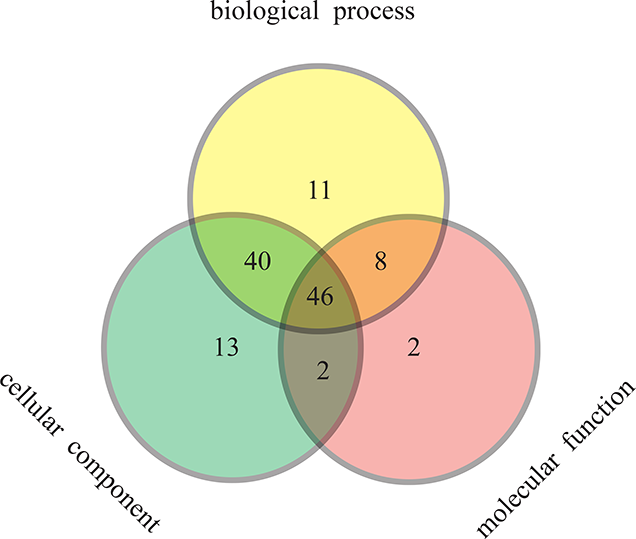
The significant difference results of ANOVA (P<0.001) in different GO categories

**Fig. 4.**
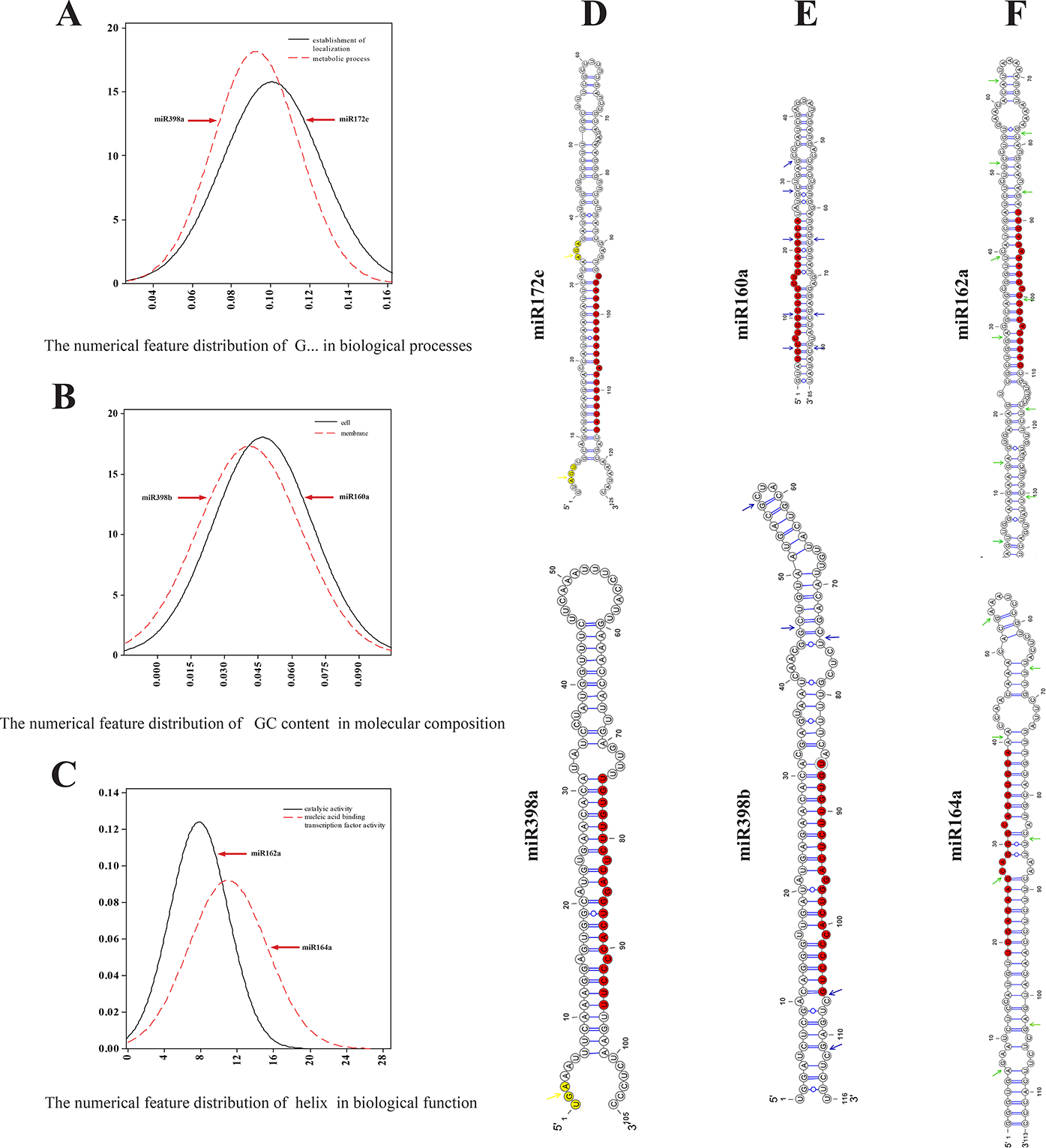
The probability density curve and the case of structure of miRNAs. A, B and C represents the distribution of numerical of different GO terms. D. The case of G… numerical feature that the yellow arrow indicates the characteristic sites. E. The case of GC content numerical feature that the blue arrow indicates the characteristic sites. F. The case of helix numerical feature that the green arrow indicates the characteristic sites. All the mature of miRNAs showed in red. The VARNA(http://varna.lri.fr/index.php?lang=en&page=home&css=varna) was used to visualize the structure of miRNAs[28].

The numerical feature GC content, as a function lable, could distinguish the ‘cell’ and ‘membrane’ in cellular component within miR398b/160a. The miR160a participated in seed germination of Lotus japonicas[33]. The case of miR162a/164a showed the numerical feature Helix could distinguish ‘catalyic activity’ and ‘nucleic acid binding transcription factor activity’ in molecular function. The evidence indicated that miR162a were experimentally verified to be the targets of nodal modulator 1-like protein gene[34]. The CUC1 and CUC2 transcripts are targeted by miR164 family, encoded by MIR164a, b and c[35]. GC content and Helix are the affect factors about the secondary structure of pre-miRNA. The more GC content and Helix in single pre-miRNA the higher energies they have. Mature miRNAs are the products of inverted repeat transcripts that are precisely cleaved by RNase III enzyme(s) in the Dicer and/or Drosha protein families[36] may need more greater energies. The previous evidences have shown that unlike other non-coding RNAs, the microRNA precursors have lower MFE (minimum free energy) than random sequences[12].

In addition, the numerical feature could not accurately describe all function of miRNAs in palnt. There are many computational predicted miRNAs need to experiment to verify and some function were unknown, because of the low quality species genome. With the development of more numerical characteristics and the next-generation sequencing technology, the miRNA biogenesis its regulatory network will be more clear in the near future.

